# Multi-scale transcriptomic integration reveals cell-type immune networks and lncRNA remodeling in Alzheimer’s disease

**DOI:** 10.64898/2026.02.08.704661

**Authors:** Jonathan Alexis Cortés Silva, Michael Gertig, Tonatiuh Pena Centeno, Susanne Burkhardt, Anna-Lena Schütz, Farahnaz Sananbenesi, André Fischer

## Abstract

Alzheimer’s disease (AD) displays pronounced regional heterogeneity, yet how transcriptional changes across brain regions converge into coordinated cellular and molecular programs remains unclear. Here, we integrated bulk and single-cell transcriptomics with network modeling to characterize gene expression remodeling across cortical and hippocampal subregions in APP/PS1-21 mice. We show that amyloid pathology follows distinct regional trajectories, with early cortical activation, delayed but robust remodeling in CA1, and a late-stage shift toward widespread transcriptional repression in the dentate gyrus. Despite these differences, cross-region analyses revealed a conserved immune activation core spanning cortical and hippocampal circuits. Network-level modeling further demonstrated that disease-associated transcriptional changes organize into immune-enriched modules that map onto specific cellular compartments, predominantly associated with microglia in cortex, astrocytes in CA1, and coordinated multi-lineage remodeling in the dentate gyrus. Notably, long noncoding RNAs were consistently embedded within disease-associated networks despite weak single-cell differential expression signals, suggesting their involvement in coordinated regulatory programs. Together, these findings link regional transcriptomic remodeling to cell-type–resolved network architecture and identify convergent immune-driven programs underlying amyloid-associated neurodegeneration.

## Introduction

Alzheimer’s disease (AD) arises on the pathological background of amyloid and Tau pathology and is characterized by progressive neurodegeneration accompanied by sustained neuroinflammatory activation that displays marked regional heterogeneity and stage-dependent dynamics (Serrano-Pozo et al., 2011; Hansen et al., 2018; Thal et al., 2002). Accumulating evidence indicates that immune responses are not merely secondary consequences of amyloid pathology, but also act as active drivers of synaptic dysfunction, neuronal vulnerability, and circuit-level degeneration (Heneka et al., 2015; Hong et al., 2016; Rajendran & Paolicelli, 2018; Leng and Edison, 2021). Consistent with this view, transcriptomic studies in human postmortem tissue and mouse models have revealed robust upregulation of glial and immune-associated gene programs, highlighting microglia and astrocytes as important mediators of AD-associated molecular remodeling (Mathys et al., 2019; Keren-Shaul et al., 2017; Grubman et al., 2019; Sala Frigerio et al., 2019).

Despite these advances, important gaps remain in understanding how transcriptional responses unfold across anatomically and functionally distinct brain regions. AD pathology exhibits pronounced regional heterogeneity, yet most transcriptomic studies focus on single brain regions or pooled tissue samples, thereby limiting the ability to resolve region-specific disease trajectories (Habib et al., 2020; Wang et al., 2016). In parallel, commonly used transcriptomic approaches provide complementary but incomplete views of disease biology. Bulk RNA sequencing enables robust detection of coordinated gene expression changes but lacks cellular resolution (Kang et al., 2011), whereas single-cell approaches offer cell-type– specific profiling but often fail to capture low-abundance regulatory transcripts and higher-order network organization due to technical sparsity and dropout effects inherent to single-cell measurements (Svensson et al., 2018; Van den Berge et al., 2020).

Integrative strategies that combine these modalities are therefore required to link regional transcriptomic remodeling with cell-type–resolved molecular architecture (Stuart et al., 2019; Argelaguet et al., 2018). Such integrative frameworks are particularly important for capturing coordinated regulatory programs that span multiple cellular compartments. Long noncoding RNAs (lncRNAs) have also emerged as important regulatory components of neuroinflammatory signaling, chromatin organization, and transcriptional coordination in neurodegenerative disease (Wan et al., 2017; Magistri et al., 2015; Qureshi & Mehler, 2012; Statello et al., 2021). However, their contribution to disease-associated transcriptional networks, particularly in a region-specific and cell-type–resolved framework, remains largely unexplored (Guennewig & Cooper, 2014).

Here, we address these challenges by integrating bulk RNA sequencing across multiple brain regions with co-expression network analysis and single-cell transcriptomic mapping in the APP/PS1-21 mouse model of amyloid pathology. By profiling the anterior cingulate cortex, hippocampal CA1 region, and dentate gyrus across disease stages, we first define region-specific transcriptional trajectories, then identify a conserved immune activation core and its network-level organization, and finally map disease-associated modules to distinct cellular compartments. This multi-scale framework provides an integrated view of how amyloid pathology reorganizes transcriptional programs across brain regions and cell populations during Alzheimer’s disease progression

## Materials and Methods

### Animals

For bulk RNA-sequencing experiments, we used tissue samples from the APP/PS1-21 transgenic mouse line B6-Tg(Thy1-APPswe; Thy1-PS1 L166P), which co-expresses human APP carrying the Swedish mutation (K670N/M671L) and human PSEN1 harboring the L166P substitution under the Thy1 promoter. This model recapitulates early and progressive amyloid pathology. Experimental cohorts consisted of heterozygous transgenic mice (APP/ PS1) and their wild-type (WT) littermates, distributed by genotype across age groups. We obtained tissue samples from animals that were 1.5 months (pre-pathology), 4 months (early amyloid deposition), and 8 months (prominent pathology) of age.

### Brain dissection and tissue processing

Mice were euthanized by cervical dislocation followed by decapitation. Brains were rapidly removed, placed on ice, and microdissected to isolate the anterior cingulate cortex (ACC), hippocampal CA1 subfield (CA1), and dentate gyrus (DG). Dissected tissue was maintained under RNase-minimizing conditions and stored appropriately until RNA extraction.

### RNA isolation

RNA isolation was performed under RNase-free conditions. Tissue was homogenized in TRIzol reagent (Thermo Fisher Scientific) with ceramic beads using an Omni Bead-Ruptor 24, followed by phase separation with chloroform. RNA was precipitated with isopropanol in the presence of GlycoBlue, washed twice with 75% ethanol, briefly air-dried, resuspended in sequencing-grade water, and incubated at 42 °C prior to DNase treatment.

### Bulk RNA-seq library preparation and sequencing

Genome-wide mRNA profiling from ACC, CA1, and DG of WT and APP/PS1 mice aged 1.5, 4, and 8 months (n = 6 per experimental group) was performed on an Illumina HiSeq 2000 platform. Paired-end libraries were prepared for CA1, whereas single-end libraries were generated for ACC and DG using the TruSeq RNA Library Preparation Kit v2. Cluster generation used TruSeq PE Cluster Kit v3-cBot-HS or TruSeq SR Cluster Kit v3-cBot-HS following Illumina standard protocols. Library quality was assessed using NanoDrop 8000, Qubit 2.0, and the RNA 6000 Nano Kit on an Agilent 2100 Bioanalyzer.

### Transcriptomic data processing and analysis

All computational analyses were performed in R (v4.4.3). Key packages included DESeq2 (v1.46.0), WGCNA (v1.73), Seurat (v5.3.0), glmGamPoi (v1.18.0), sctransform (v0.4.2), clusterProfiler (v4.14.6), org.Mm.eg.db (v3.20.0), enrichplot (v1.26.6), biomaRt (v2.62.1), and ggplot2 (v3.5.2).

### Bulk RNA-seq differential expression analysis

Bulk RNA-seq analyses were performed independently for ACC, CA1, and DG using an identical pipeline to ensure methodological consistency across anatomical regions and time points. Gene-level count matrices were generated from aligned reads and integrated with sample metadata. Differential expression analyses were conducted using DESeq2 on integer counts. Samples were modeled using a composite experimental factor encoding genotype and age (condition_time = genotype_timepoint), with design formula ∼ condition_time. Genotype-matched contrasts were tested independently at each disease stage (APP vs WT at 1.5, 4, and 8 months).

Genes were considered significantly differentially expressed using a Benjamini–Hochberg FDR < 0.05 and an absolute log2 fold-change ≥ 0.26. A light abundance filter was applied prior to inference, retaining genes with a mean normalized count > 10 across samples. Variance-stabilized expression matrices were computed using DESeq2 vst() and used for PCA, clustering, and network construction. Log2 fold changes were reported without shrinkage.

### lncRNA differential expression analysis

Long noncoding RNAs were defined from GTF annotations for Mus musculus GRCm39 (release 105) by selecting genes annotated with gene_biotype = “lncRNA”. Count matrices were subset to lncRNAs and analyzed using the same DESeq2 design (∼ condition_time), contrasts (APP vs WT at 1.5/4/8 months), abundance filter (mean normalized count > 10), and significance thresholds (FDR < 0.05, |log2FC| ≥ 0.26) applied to protein-coding genes. Variance-stabilized lncRNA matrices were used for PCA and visualization.

### Weighted gene co-expression network analysis (WGCNA)

Co-expression networks were constructed independently for ACC/PFC, CA1, and DG using DESeq2 VST expression values. For each region, genes were ranked by variance and the top 5,000 most variable genes were used as input. Expression matrices (samples × genes) were screened using goodSamplesGenes. Signed networks were computed using robust biweight midcorrelation (bicor) with maxPOutliers = 0.05.

Soft-thresholding powers were evaluated using pickSoftThreshold (candidate powers 1–10 and 12–30, step size 2), targeting a signed scale-free topology fit index R^2^ ≥ 0.80; when not achieved, the power yielding the highest R^2^ was selected. Final selected powers were 9 (ACC/PFC), 12 (CA1), and 12 (DG). Modules were identified using blockwiseModules with minModuleSize = 40, deepSplit = 2, mergeCutHeight = 0.20, reassignThreshold = 0, and PAM refinement respecting dendrogram structure. Module eigengenes were ordered using orderMEs (excluding the grey module) and eigengene orientation was standardized to maintain consistent directionality with respect to experimental traits (Condition and Time). Module assignments were exported for integration with single-cell pseudobulk results.

### GO enrichment analysis

Gene Ontology enrichment analyses were performed using clusterProfiler (v4.14.6) with mouse annotation from org.Mm.eg.db (v3.20.0) and visualization using enrichplot (v1.26.6). The background universe was defined per region using a fixed background gene list derived from the network analysis (GO_background_geneList.csv), comprising 4855 genes (ACC/ PFC), 4866 genes (CA1), and 4862 genes (DG) (Ensembl gene IDs). Multiple-testing correction used the Benjamini–Hochberg procedure.

### Single-cell RNA-seq preprocessing, integration, and pseudobulk analysis

Public scRNA-seq datasets were obtained from GEO (GSE221959, GSE173242, GSE198027, GSE221702) and processed using a unified Seurat (v5.3.0) workflow. Raw 10x Genomics matrices (matrix.mtx, barcodes.tsv, features.tsv) were loaded per sample, duplicate gene names were disambiguated, and cell barcodes were prefixed with sample identifiers. Quality control metrics included detected genes, UMI counts, mitochondrial fraction (^mt-), and ribosomal fraction (^Rpl, ^Rps). Cells were filtered using sample-specific thresholds based on median ± 3 MAD for gene and UMI counts, with high mitochondrial and ribosomal outliers removed using the 99.5th percentile cutoff; samples retaining fewer than 50 cells were excluded. Cortex datasets were downsampled to a maximum of 3,000 cells per sample and 40,000 total cells prior to integration.

Normalization was performed using SCTransform (3,000 variable features) with glmGamPoi acceleration when available. SCT-based integration used SelectIntegrationFeatures (2,000 features), PrepSCTIntegration, FindIntegrationAnchors, and IntegrateData, performed separately for cortex and hippocampus to minimize region-driven batch effects. Dimensionality reduction used PCA, with the first 30 PCs used for neighbor graph construction, UMAP embedding, and clustering. Clustering was performed with resolution 0.6. Cell types were assigned using canonical marker gene panels by computing average marker expression per cluster and assigning the highest-scoring label, followed by marker-based validation.

For pseudobulk analysis, raw counts were aggregated per SampleID × CellType by summing counts across cells. Only strata with at least 30 cells and representing both conditions (TG and Control) were retained. Differential expression was performed using DESeq2 with design ∼ Condition, testing TG vs Control independently for each mouse model and cell type.

### Integration of pseudobulk signatures with bulk WGCNA modules

Pseudobulk DE results were merged with bulk WGCNA module assignments using gene symbols. For each mouse model and cell type, enrichment of differentially expressed genes (FDR < 0.05) within each WGCNA module was assessed using Fisher’s exact test based on a 2×2 contingency table of DEG vs non-DEG genes inside and outside the module. P-values were adjusted for multiple testing within each mouse model and cell type using FDR correction. Enrichment results were summarized and visualized using dot plots encoding DEG fraction and enrichment significance.

To investigate how amyloid pathology reshapes transcriptional programs across anatomically distinct cortical and hippocampal regions, we performed bulk RNA sequencing across multiple disease stages. This analysis was designed to define region-specific transcriptional trajectories and establish a foundation for subsequent network-level and cell-type resolved analyses.

## Results

### Regional transcriptional remodeling reveals distinct spatiotemporal trajectories across cortical and hippocampal regions

APP/PS1-21 mice develop an early-onset cerebral amyloidosis driven by neuronally expressed human APP (Swedish mutation) and PS1 (L166P). Amyloid deposition begins first in the neocortex around 1.5 months, with rapidly increasing cortical plaque load and prominent plaque-associated dystrophic neurites. In parallel, a progressive neuroinflammatory response emerges (Radde et al., 2006). By 4 months, amyloid pathology is well established in cortex and becomes evident in hippocampus, which is accompanied by increasing microglial/astrocytic activation and plaque-associated synaptic pathology. By 8 months, plaque burden is high in cortex and more pronounced in hippocampus, with strong gliosis and plaque-proximal structural synaptic deficits. At this stage, hippocampal CA1 synaptic plasticity becomes markedly impaired as indicated by LTP deficits and learning impairments (Gengler et al., 2010).

Since a systematic transcriptome analysis of APP/PS1-21 mice is lacking, we performed RNA sequencing in the anterior cingulate cortex (ACC), hippocampal CA1 region, and dentate gyrus (DG) of APP/PS1-21 mice at pre-pathological (1.5 months), early amyloid (4 months), and advanced amyloid pathology (8 months) stages. This experimental design enabled a systematic comparison of transcriptional dynamics across anatomically and functionally distinct brain regions during disease progression. Volcano plot analysis of protein-coding gene expression revealed marked regional differences in both the magnitude and direction of transcriptional remodeling (Figure 1).

**Figure 1.**
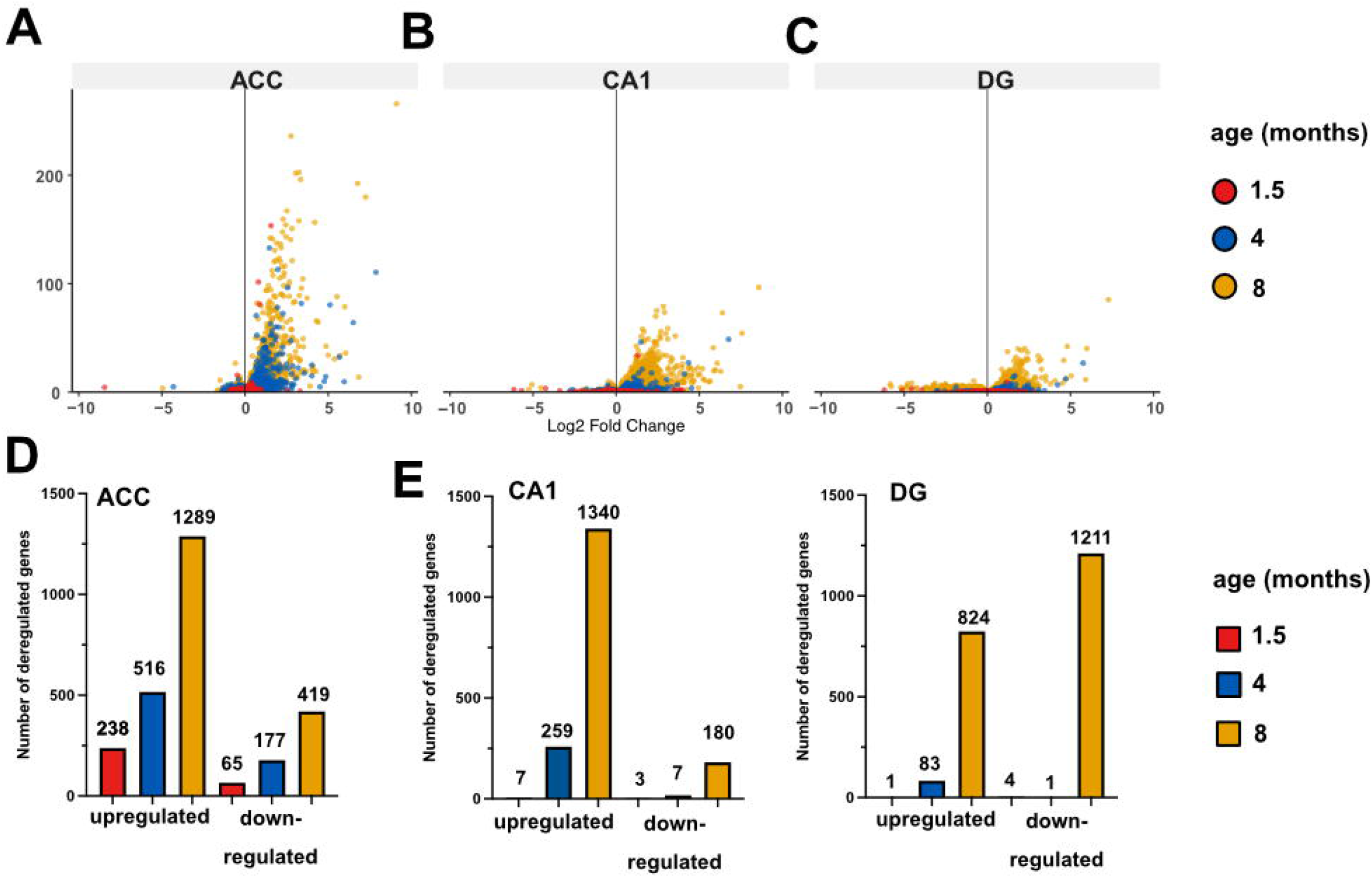
RNA-seq reveals distinct spatiotemporal transcriptional trajectories across ACC, CA1, and DG during amyloid pathology progression in APP/PS1-21 mice. A. Volcano plots showing protein-coding genes that are differentially expressed in the ACC between control and APP/PS1-21 mice at 1.5, 4, and 8 months of age. The same analysis was performed for the hippocampal CA1 region (B) and the DG (C). For all brain regions, time points and genotypes 6 mice/groups were analyzed. FDR (padj) < 0.05 and |log2FC| ≥ 0.26 were applied to all panels. D. Bar chart summarizing the number of significantly upregulated and downregulated protein-coding genes in the ACC, the CA1 region (E) and the DG (F).

In the ACC, widespread transcriptional changes were already observed in 1.5-month-old APP/PS1-21 mice compared with control littermates. Transcriptional dysregulation gradually increased at 4 and 8 months of age, with more genes being upregulated than downregulated (Fig. 1A, D).

In the CA1 region, only minimal transcriptional perturbations were observed in 1.5-month-old mice. At 4 months, 259 upregulated genes were detected, whereas only 7 gene**s** showed decreased expression. At 8 months, 1,340 genes were significantly upregulated, while 180 genes showed significantly decreased expression (Fig. 1B, E). By comparison, the DG exhibited only 1 upregulated and 4 downregulated genes at 1.5 months, and 83 upregulated and 1 downregulated gene at 4 months. At 8 months, we detected 824 increased genes and 1211 decreased genes (Fig. 1C, F).

Together, these results demonstrate that amyloid-driven transcriptional remodeling follows highly distinct spatiotemporal trajectories across cortical and hippocampal circuits. These trajectories align well with the reported development of amyloid pathology in APP/PS1-21 mice, namely the fact that cortical regions are affected first, with amyloid pathology detectable as early as 1.5 months of age, a time point when hippocampal tissues are still largely intact (Radde, 2006).

### Cross-region convergence reveals a shared immune activation core and network-level integration of disease-associated programs

To identify transcriptional programs conserved across cortical and hippocampal circuits, we analyzed the overlap of differentially expressed protein-coding genes across the ACC, CA1, and DG. For each region, differentially expressed genes (DEGs) detected at 1.5, 4, and 8 months were aggregated to generate region-level disease-associated gene sets. Intersection analysis revealed a substantial disease-associated gene core shared across all three brain regions (551 protein-coding genes, Fig. 2A), indicating the presence of conserved transcriptional responses that extend across multiple stages of pathogenesis. At the same time, region-specific DEG subsets were retained, suggesting the coexistence of shared and anatomically selective molecular programs.

**Figure 2.**
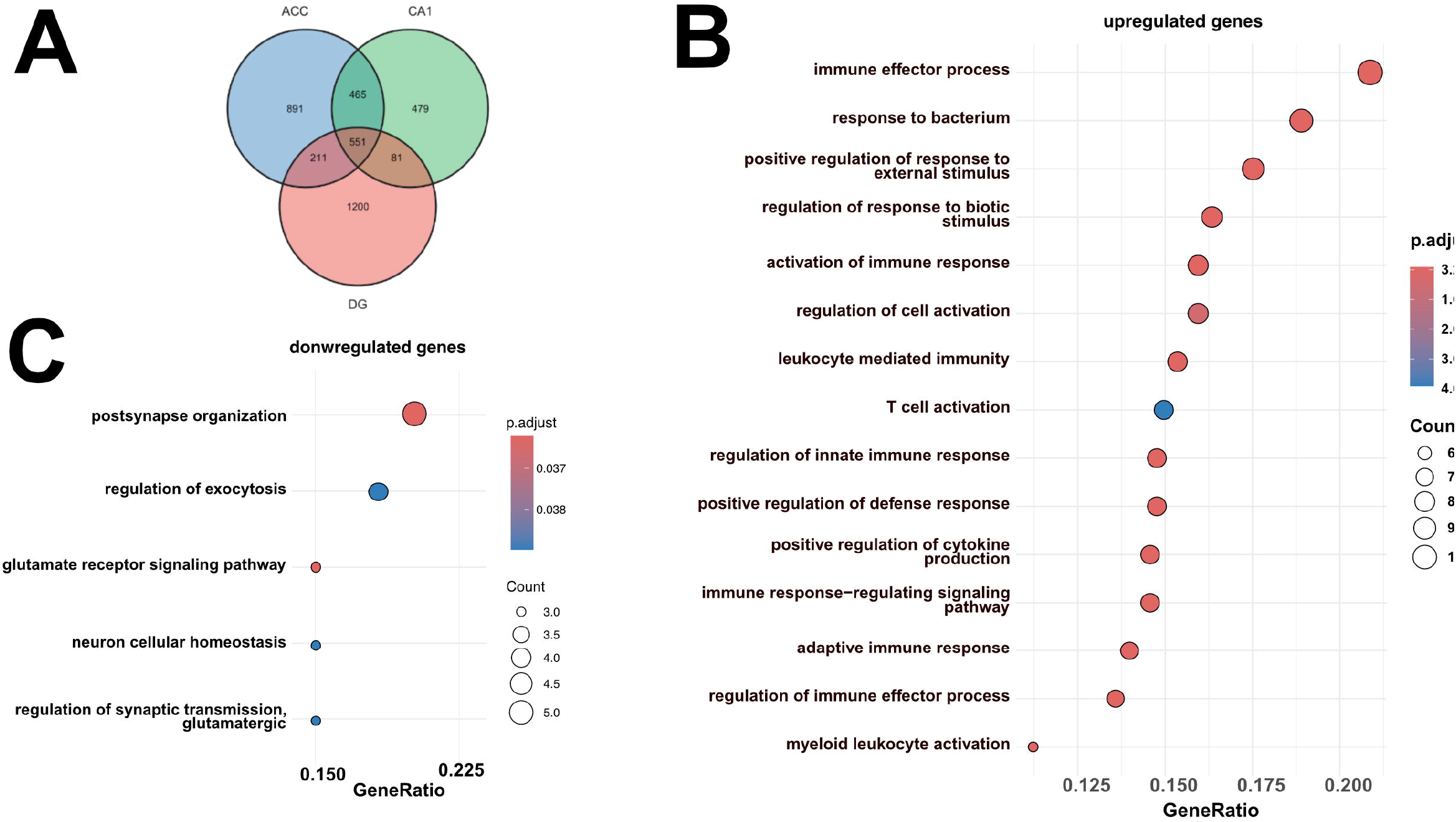
Cross-regional overlap across disease stages reveals a shared immune-related protein-coding signature. A. Venn diagram showing overlap of protein-coding differentially expressed genes (DEGs) across ACC, CA1, and DG, computed by using all analyzed time points (1.5, 4, and 8 months) for each region (DEGs defined as the union across time points using the DESeq2 model ∼condition_time and an abundance filter of mean normalized counts >10).Within each region, DEGs were defined using identical statistical thresholds (FDR < 0.05 and |log2FC| > 0.26). Under this definition, 551 protein-coding genes were shared across all three regions (ACC–CA1–DG), with pairwise overlaps of 1016 (ACC–CA1), 762 (ACC–DG), and 632 (CA1–DG). The total numbers of unique protein-coding DEGs across time were 2118 in ACC, 1576 in CA1, and 2043 in DG. B. Dot blot showing GO term (Biological Process) enrichment analysis of the cross-region shared protein-coding gene set of upregulated genes (n = 508). C. Dot blot showing GO term (Biological Process) enrichment analysis of the cross-region shared protein-coding gene set of downregulated genes (n = 22).

Gene ontology (GO) analysis of the shared protein-coding transcripts showed that these were consistently upregulated in APP/PS1-21 mice across brain regions and time points were almost exclusively enriched for processes linked to inflammatory pathways such as “immune effector processes”, “response to bacterium”, “regulation of innate immune response”, “T cell activation” or “adaptive immune response” (Fig. 2B). GO term analysis of the shared downregulated genes revealed processes such as “postsynapse organization”, “Regulation of Exocytosis”, “glutamate receptor signaling pathways” or “neuron cellular homeostasis” neuronal and synaptic processes, including glutamatergic signaling and postsynaptic organization (Fig. 2C), suggesting convergent impairment of synaptic-related pathways across anatomically distinct brain regions during disease progression. These data are in agreement with previous studies suggesting that, in amyloid pathology models, transcriptional changes precede the dysregulation of genes linked to synaptic and neuronal plasticity (Matarin et al., 2015) and place amyloid deposition–mediated neuroinflammation at the top of a disease cascade that eventually leads to synaptic disturbances.

To determine whether disease-associated transcriptional changes were organized into coherent gene networks, we next performed weighted gene co-expression network analysis (WGCNA). Here, we constructed region-specific co-expression networks using variance-stabilized expression (DESeq2 VST) and the top 5,000 most variable genes per region as input. Module–trait correlation analysis identified multiple disease-associated modules significantly linked to genotype and/or disease progression in ACC (Fig 3A), CA1 (Fig 3B). and DG (Fig. 3C). For downstream analyses, we focused on selected representative modules showing the strongest disease associations and robust functional enrichment.

**Figure 3.**
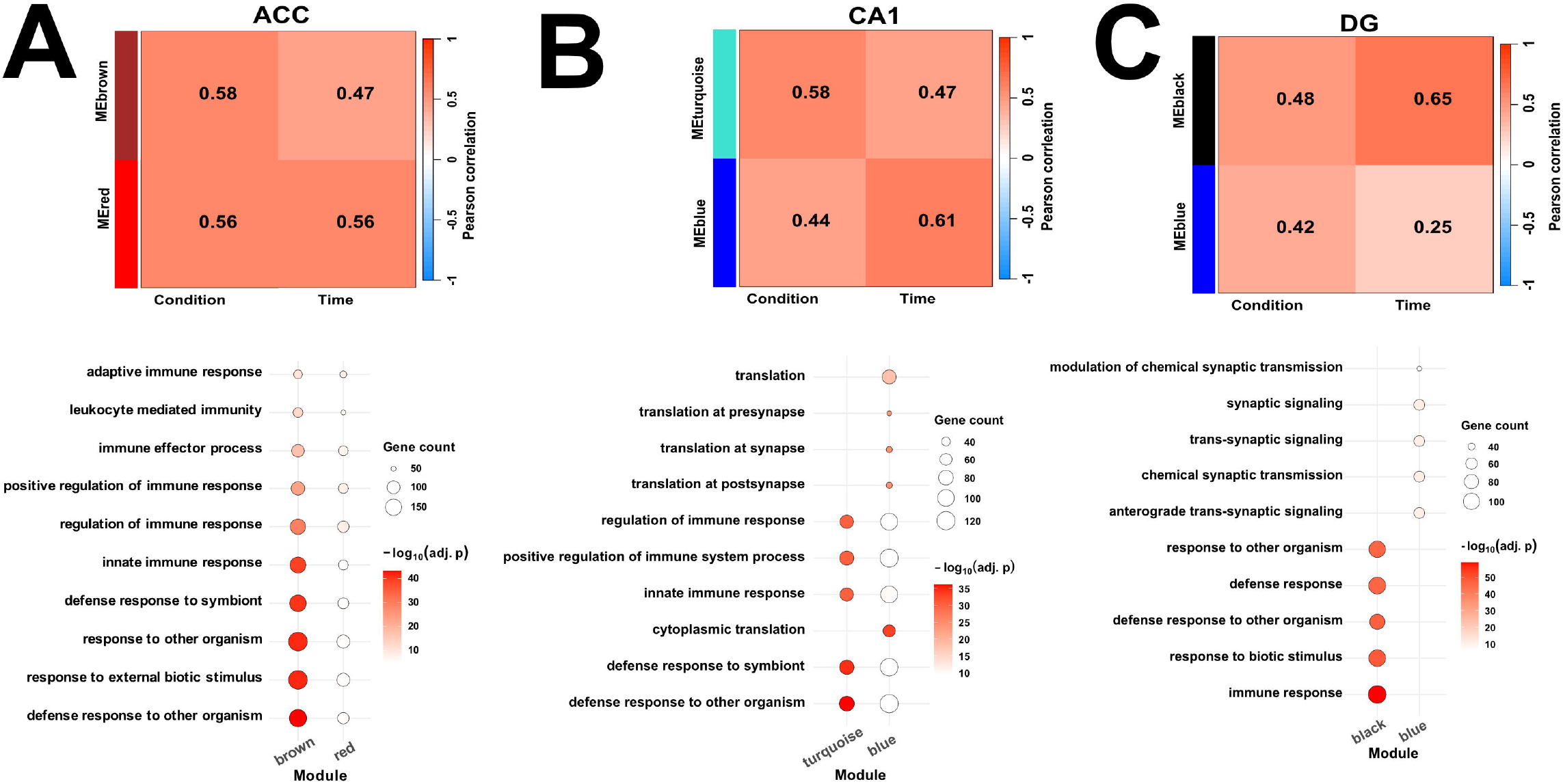
Cross-region network-level analysis across disease stages identifies a conserved immune-enriched program. A. Upper panel: WGCNA module–trait correlation heatmap from region-specific co-expression networks constructed from the expressed transcriptome (top 5000 most variable genes per region), integrating protein-coding and lncRNA transcripts. Modules shown were prioritized as disease-associated based on significant correlations with genotype and/or age. Lower panel: GO Biological Process enrichment analysis of disease-associated WGCNA modules. B. The same analysis as described under (A) was performed for the CA1 region and the DG (C).

Functional annotation of prioritized modules revealed strong enrichment for immune and inflammatory biological processes across all regions. In the ACC, disease-associated modules were enriched for pathways related to innate immune activation and inflammatory signaling (Fig. 3A). Similarly, CA1 (Fig. 3B) and DG (Fig. 3C) modules displayed enrichment for immune-related and stress-responsive biological processes along with processes linked to synaptic function, which was particularly prominent in the DG, most likely reflecting the higher number of down-regulated genes in 8 months old mice that were mainly linked to synaptic processes. Together, these results suggest that immune-driven transcriptional programs represent a dominant organizing principle of amyloid-associated network architecture across cortical and hippocampal regions.

Collectively, these data further suggest that amyloid pathology engages a conserved immune activation core that emerges across anatomically distinct brain regions and becomes robustly detectable at the gene level at advanced disease stages, while simultaneously reorganizing region-specific transcriptional networks that integrate both protein-coding genes and regulatory noncoding transcripts.

### Single-cell integration localizes disease-associated networks to distinct cellular compartments

To resolve the cellular origins of the disease-associated transcriptional networks identified in our study, we integrated WGCNA module information with pseudobulk differential expression profiles derived from single-cell RNA sequencing datasets spanning multiple AD mouse models for amyloid pathology. This integrative approach enabled systematic mapping of region-specific co-expression modules onto defined brain cell populations, providing cellular context to bulk transcriptomic patterns.

In the ACC, disease-associated modules showed strong enrichment for microglial transcriptional signatures (Fig. 4A). This is in agreement with our observation that cortical gene-expression changes in our model system mainly reflect immune and innate response programs. A weaker yet significant contribution was observed for oligodendrocyte precursor.

**Figure 4.**
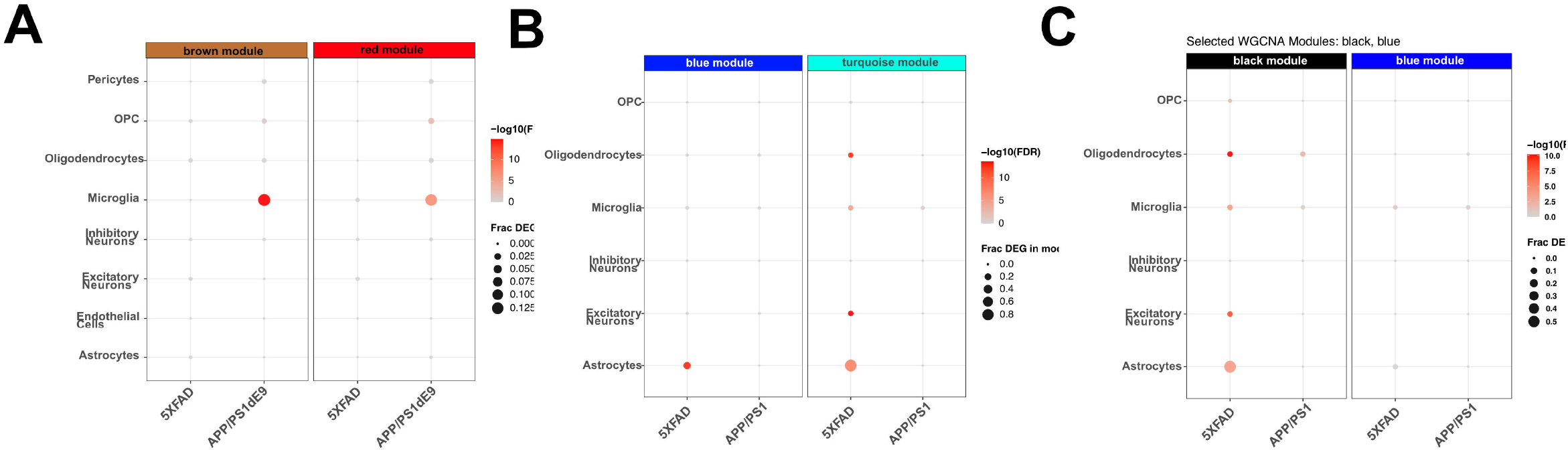
Pseudobulk single-cell integration maps disease-associated WGCNA modules to specific cell populations across regions. A. Dot blot showing disease-associated WGCNA modules from the ACC that were integrated with pseudobulk single-cell differential expression using Fisher’s exact tests for overlap between each module gene set and each cell type’s DEG list within each mouse model, using module gene sets derived from bulk WGCNA networks. Enrichment results are reported as odds ratio (OR) with multiple-testing correction (padj/FDR). Dot plots summarize enrichment across cell types and mouse models. Significant enrichment was observed in microglia for both modules (brown: FDR 1.37×10^−15^; red: FDR 1.10×10^−7^), with an additional significant signal for the red module in OPCs (FDR 1.98×10^−3^). Effect sizes were consistent with these signals (median OR: brown–microglia 3.89; red–microglia 2.91; red–OPC 9.70). Maximum module coverage among significant results was 0.133 (brown– microglia) and 0.125 (red–microglia). B. A similar analysis as shown in (A) was performed for the hippocampal CA1 region. The turquoise module showed strong enrichment across multiple compartments, including excitatory neurons (FDR 2.80×10^−14^), astrocytes (FDR 5.56×10^−8^), oligodendrocytes (FDR 3.17×10^−13^), and microglia (FDR 1.95×10^−5^). The blue module showed significant enrichment in astrocytes (FDR 1.59×10^−13^). Estimated effect sizes were large for turquoise– astrocyte and turquoise–oligodendrocyte overlaps (median OR 109.7 and 70.1, respectively), with high fractional coverage for turquoise–astrocyte (max frac_DEG_in_module 0.889). C. A similar analysis as shown in (A) was performed for the DG. Significant enrichment was dominated by the black module, with strongest signals in oligodendrocytes (FDR 6.18×10^−11^) and excitatory neurons (FDR 4.59×10^−9^), and additional significant enrichment in astrocytes (FDR 1.16×10^−4^), microglia (FDR 1.83×10^−4^), and OPCs (FDR 1.39×10^−2^). Reported effect sizes included high median ORs for black– oligodendrocyte (33.3), black–astrocyte (22.5), and black–excitatory (17.8); black–OPC returned OR=Inf in the output (typically reflecting a zero cell in the contingency table).

In contrast, in the hippocampal CA1 region the module showing the strongest disease association was enriched for astrocyte-associated transcriptional signatures, with additional, yet less significant contributions from excitatory neurons and oligodendrocytes (Fig. 4B).

Gene-expression changes in the DG showed enrichment across multiple cell populations, including astrocytes, oligodendrocytes, excitatory neurons, and microglia. Among these, astrocytes accounted for the largest fraction of module-associated transcriptional changes (Fig. 4C).

Together, these findings support the view that region-specific transcriptional networks identified by bulk RNA sequencing correspond to distinct cellular drivers across brain regions. While cortical changes in gene-expression can be attributed primarily to microglial activation, hippocampal CA1 gene-expression changes are strongly associated with astrocyte-dependent processes, whereas DG remodeling reflects coordinated multi-cellular responses involving astrocytes, oligodendrocytes, neurons, and microglia.

### Regional changes in lncRNA expression

Most therapeutic and mechanistic work has historically focused on protein-coding genes (∼1.5% of the genome), reflecting the fact that proteins are direct drug targets. However, the majority of genomic output is non-coding, including long non-coding RNAs (lncRNAs), generally defined as >300-nt transcripts without protein-coding potential (Statello et al., 2021; Bridges et al., 2021). Rather than representing transcriptional “noise,” lncRNAs are increasingly recognized as functional regulators in many biological contexts (Ransohoff et al., 2018; Quinn & Chang, 2016). Because lncRNAs are often highly tissue- and cell type– specific and are frequently enriched in the brain (Mercer et al., 2012), they are well positioned to shape region- and cell-specific responses relevant to brain disorders (Qureshi & Mehler, 2012; Riva et al., 2016; Wan & Zhuo, 2017; Briggs et al., 2015; Liu et al., 2021).

Therefore, we also analyzed differential expression of lncRNAs. Compared with the changes observed for protein-coding transcripts (see Fig. 1), fewer lncRNAs were differentially expressed overall. However, the temporal trajectories broadly paralleled those observed for protein-coding transcripts.

In the ACC, lncRNA differential expression progressively expanded across disease stages, with the strongest response observed at 8 months (Fig. 5A, D). Specifically, the ACC showed a gradual increase from 2 upregulated lncRNAs at 1.5 months, to 3 at 4 months, and to 32 at 8 months of age. No downregulated lncRNAs were detected at 1.5 months, whereas 3 and 19 lncRNAs were downregulated at 4 and 8 months of age, respectively (Fig. 5A, D). In the hippocampal CA1 region, no downregulated lncRNAs were detected at any time point. One lncRNA was increased at 1.5 and 4 months of age, and 19 were significantly elevated at 8 months of age (Fig. 5B, E). In the DG, no differentially expressed lncRNAs were detected at 1.5 months of age, and one upregulated and one downregulated lncRNA were observed at 4 months of age. At 8 months of age, 37 lncRNAs were decreased and 15 were increased (Fig. 5C, F).

**Figure 5.**
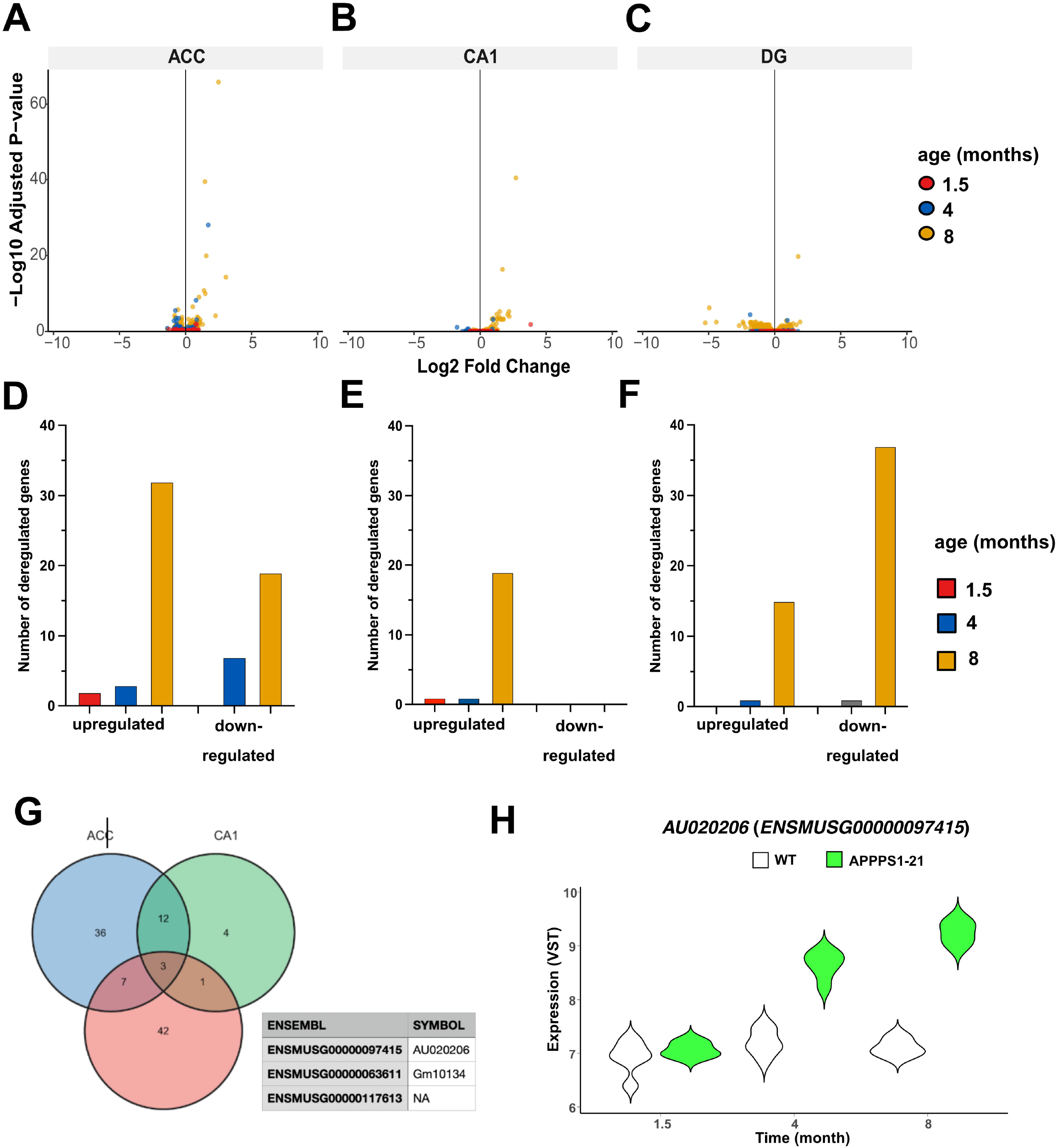
RNA-seq reveals distinct spatiotemporal changes in lncRNA expression across ACC, CA1, and DG during amyloid pathology progression in APP/PS1-21 mice. A. Volcano plots showing lncRNAs that are differentially expressed in the ACC between control and APP/PS1-21 mice at 1.5, 4, and 8 months of age. The same analysis was performed for the hippocampal CA1 region (B) and the DG (C). For all brain regions, time points and genotypes 6 mice/ groups were analyzed. FDR (padj) < 0.05 and |log2FC| ≥ 0.26 were applied to all panels. D. Bar chart summarizing the number of significantly upregulated and downregulated lncRNAs in the ACC, the CA1 region (E) and the DG (F). G. Venn diagram showing overlap of differentially expressed lncRNA across ACC, CA1, and DG. Only three lncRNAs were shared across all regions (See lower panel): ENSMUSG00000097415 (AU020206), ENSMUSG00000117613, and ENSMUSG00000063611. H. Violin plot showing the expression of AU020206 in WT and APP/PS1-21 mice. AU020206 module membership reflects co-expression network structure and does not imply single-gene differential expression in individual contrasts.

When we applied the same cross-region convergence approach used to study protein-coding transcripts (see Fig. 1), we observed that differentially expressed long non-coding RNAs showed limited overlap across regions, with only three transcripts shared among the ACC, CA1, and DG (Fig. 5G). Among these shared lncRNAs, *AU020206* was of particular interest because it has a human homolog and was similarly regulated across the ACC, CA1, and DG regions (Fig. 5H).

## Discussion

### Regional heterogeneity and temporal divergence of transcriptional remodeling in amyloid pathology

Our results demonstrate that amyloid-driven transcriptional remodeling follows distinct spatiotemporal trajectories across cortical and hippocampal circuits. The anterior cingulate cortex exhibited early and sustained transcriptional activation, whereas CA1 showed delayed but robust late-stage remodeling, and the dentate gyrus transitioned into a state dominated by widespread transcriptional repression. These findings are consistent with the concept that Alzheimer’s disease does not progress uniformly across the brain but instead affects regions with different temporal dynamics and vulnerabilities. This regional divergence aligns with previous observations that Alzheimer’s disease progression is shaped by circuit architecture, cellular composition, and local inflammatory environments (Braak and Braak, 1991; Roussarie et al., 2020).

Early cortical transcriptional activation observed in the ACC may reflect heightened sensitivity of frontal and association cortices to soluble amyloid species and early microglial activation. Human transcriptomic studies have similarly reported early immune and glial signatures in cortical regions prior to extensive neuronal loss (Mathys et al., 2019; Morabito et al., 2021). While our data do not directly establish causality, they support a model in which early cortical immune activation accompanies the initial stages of amyloid-associated molecular remodeling. In contrast, the delayed response in CA1 aligns with reports that hippocampal pyramidal neuron vulnerability often emerges after initial cortical inflammatory changes, suggesting a staged propagation of molecular pathology across connected circuits (Habib et al., 2020).

The pronounced late-stage transcriptional repression observed in the dentate gyrus represents a particularly notable feature of our dataset. This pattern may reflect advanced network dysfunction, cellular exhaustion, or reduced transcriptional plasticity in this region, which plays a critical role in hippocampal neurogenesis and memory encoding. Previous studies have reported impaired neurogenic capacity and metabolic dysregulation in the dentate gyrus during advanced amyloid pathology, supporting the idea that this region follows a distinct degenerative trajectory compared to CA1 and cortical areas (Mu & Gage, 2011; Moreno-Jiménez et al., 2019). However, further work will be required to determine the mechanistic basis of this late-stage transcriptional repression.

### Conserved immune activation as a core transcriptional signature across brain regions

Despite pronounced regional heterogeneity, our cross-region analyses identified a conserved immune activation core shared across cortical and hippocampal circuits during disease progression. Genes involved in innate immune signaling, leukocyte-mediated immunity, and host-defense responses were consistently upregulated across ACC, CA1, and DG, indicating that neuroimmune activation represents a common molecular response across anatomically distinct regions. This finding reinforces the central role of neuroinflammation as a unifying feature of Alzheimer’s disease pathophysiology (Heneka et al., 2015; Leng and Edison, 2021). Importantly, this shared immune signature emerged from integrating transcriptional changes across multiple disease stages, suggesting temporal robustness of this response.

Microglial activation programs, including disease-associated microglia (DAM) signatures, have been repeatedly reported in both mouse models and human Alzheimer’s disease tissue and are thought to reflect adaptive, yet potentially maladaptive, responses to amyloid accumulation (Keren-Shaul et al., 2017; Deczkowska et al., 2018). Our results are consistent with the presence of a widespread neuroimmune response that extends beyond local circuit-specific effects. At the same time, the relatively limited overlap observed for downregulated neuronal and synaptic genes across regions suggests that neuronal dysfunction may follow more region-specific trajectories.

This dissociation between globally conserved immune activation and regionally heterogeneous neuronal remodeling highlights the importance of integrating immune-centered and neuron-centered perspectives when interpreting transcriptomic changes in neurodegenerative disease.

### Network-level organization of immune-driven transcriptional remodeling

While differential expression analyses identify individual disease-associated genes, they do not capture the coordinated structure of transcriptional regulation underlying complex biological processes. By applying weighted gene co-expression network analysis (WGCNA), we demonstrate that amyloid-associated transcriptional remodeling is organized into coherent gene modules that reflect coordinated immune and inflammatory programs across brain regions. Network-based approaches are increasingly recognized as essential for understanding systems-level organization of neurodegenerative disease biology (Zhang et al., 2013; Mostafavi et al., 2018; Allen et al., 2018).

Across ACC, CA1, and DG, disease-associated modules were consistently enriched for immune and inflammatory biological processes, including innate immune signaling, host-defense responses, and glial activation pathways. Similar immune-enriched network modules have been reported in human postmortem brain tissue and multiple AD mouse models, where co-expression networks correlate strongly with pathology and cognitive decline (Zhang et al., 2013; Mostafavi et al., 2018; Johnson et al., 2020). Our findings extend these observations by showing that immune-centered network architecture is preserved across anatomically distinct cortical and hippocampal regions.

Importantly, WGCNA enabled identification of region-specific network organization despite the presence of a shared immune core. Cortical modules in the ACC showed strong immune enrichment associated with early transcriptional activation, whereas CA1 and DG modules reflected region-dependent network restructuring at later disease stages. These observations support a model in which a broadly shared molecular response is subsequently reshaped by local cellular environments and circuit-specific vulnerabilities (Allen et al., 2018).

Network-level modeling also provides a framework to integrate regulatory elements that may not exhibit strong single-gene differential expression signals. The consistent embedding of selected lncRNAs within disease-associated modules across regions supports the idea that regulatory transcripts participate in coordinated transcriptional programs rather than acting as isolated effectors. Recent systems-level studies similarly highlight the importance of regulatory network hubs and noncoding transcripts in shaping disease-associated transcriptional states (de Jager et al., 2018; Statello et al., 2021).

Together, these results indicate that amyloid pathology reorganizes brain transcriptomes into immune-dominated co-expression networks whose regional architecture reflects both conserved inflammatory signaling and local circuit-specific modulation. This network-centric perspective links conserved immune activation with region-specific cellular responses and supports integration of bulk and single-cell transcriptomic data within a unified systems-level framework.

### Cell-type–specific drivers of network remodeling across brain regions

By integrating bulk-derived co-expression networks with pseudobulk single-cell transcriptomic profiles, we were able to assign disease-associated transcriptional programs to specific cellular compartments. This cell-type–resolved mapping revealed pronounced regional differences in the cellular drivers of network remodeling, providing mechanistic context for the heterogeneous transcriptional trajectories observed across cortical and hippocampal regions.

In the anterior cingulate cortex, disease-associated network modules were dominated by microglial transcriptional signatures, consistent with the early and sustained immune activation observed at the bulk transcriptome level. Microglia rapidly respond to amyloid deposition through activation of innate immune pathways, phagocytic programs, and inflammatory signaling cascades (Keren-Shaul et al., 2017; Deczkowska et al., 2018). Single-cell and spatial transcriptomic studies further demonstrate that microglial activation represents one of the earliest and most robust cellular responses to amyloid pathology (Zhou et al., 2020). The strong microglial enrichment of cortical networks in our data therefore supports a model in which early cortical transcriptional remodeling is closely linked to immune surveillance and inflammatory responses.

In contrast, hippocampal CA1 networks showed prominent astrocytic contributions. Astrocytes play key roles in metabolic support, synaptic regulation, and modulation of neuroinflammatory signaling, and disease-associated astrocyte states have increasingly been implicated in neurodegeneration (Habib et al., 2020; Liddelow et al., 2017). Recent transcriptomic studies suggest that reactive astrocyte programs are strongly associated with synaptic dysfunction and neuronal stress in Alzheimer’s disease (Escartin et al., 2021). The astrocytic enrichment observed in CA1 modules is therefore consistent with a potential role for astrocyte-mediated regulation of neuronal homeostasis during later disease stages.

The dentate gyrus displayed a more heterogeneous cellular profile, with disease-associated networks incorporating contributions from astrocytes, oligodendrocytes, excitatory neurons, and microglia. This multi-lineage involvement aligns with the pronounced late-stage transcriptional repression and high DEG burden observed in this region. The dentate gyrus is particularly sensitive to disruptions in cellular plasticity and neurogenic processes, and coordinated dysfunction across multiple cell types may reflect widespread network instability during advanced amyloid pathology (Mu and Gage, 2011; Moreno-Jiménez et al., 2019). Multi-cellular transcriptional convergence has also been reported in advanced neurodegenerative stages, suggesting progressive breakdown of cell-type-specific homeostatic programs (Mathys et al., 2019).

Together, these findings indicate that amyloid-associated transcriptional networks are not uniformly executed across cell populations but instead reflect region-dependent cellular drivers. While cortical remodeling appears primarily associated with microglial activation, hippocampal CA1 networks show strong astrocytic involvement, and dentate gyrus remodeling reflects coordinated multi-cellular responses. This cellular stratification provides a mechanistic bridge linking regional transcriptomic heterogeneity with distinct cellular response programs during disease progression.

### Network-level integration reveals regulatory involvement of lncRNAs in amyloid-associated transcriptional programs

Although long noncoding RNAs represented a relatively small fraction of differentially expressed transcripts compared to protein-coding genes, our network-based analyses indicate that selected lncRNAs participate in coordinated disease-associated transcriptional programs. In particular, AU020206 was consistently embedded within disease-associated co-expression modules across ACC, CA1, and dentate gyrus, despite exhibiting limited and regionally variable differential expression at the single-gene level. This observation highlights the value of network-level approaches for identifying regulatory components that may not be captured by conventional differential expression analyses.

Increasing evidence suggests that lncRNAs contribute to neurodegenerative disease through regulation of chromatin accessibility, transcriptional coordination, and immune signaling pathways (Wan et al., 2017; Qureshi & Mehler, 2012). Genome-wide studies have further shown that many lncRNAs act as scaffolds or modulators of gene regulatory networks rather than as strongly differentially expressed transcripts (Ransohoff et al., 2018; Statello et al., 2021). In the context of Alzheimer’s disease, several lncRNAs have been implicated in amyloid processing, microglial activation, and inflammatory regulation (Magistri et al., 2015; Mathys et al., 2017).

The consistent network embedding of AU020206 across brain regions suggests that this transcript may function as a regulatory component within immune-enriched transcriptional modules. However, our data are correlative and do not establish a causal role for this lncRNA in disease progression. Future experimental studies will be required to determine its functional relevance.

More broadly, these results support a model in which noncoding transcripts contribute to Alzheimer’s disease not as isolated molecular markers but as components of distributed regulatory systems that coordinate large-scale transcriptional responses. Integrating lncRNAs into co-expression network frameworks may therefore represent a promising strategy for uncovering additional regulatory layers of disease biology.

### Limitations

While this study provides a multi-scale framework to characterize amyloid-associated transcriptional remodeling, several limitations inherent to modeling human Alzheimer’s disease in experimental animals should be considered. Although the APP/PS1-21 mouse model recapitulates key features of amyloid pathology and robust neuroinflammatory activation, it does not fully capture the complexity of sporadic late-onset Alzheimer’s disease, including tau pathology, vascular contributions, metabolic alterations, and age-related comorbidities. Accordingly, extrapolation of region-specific and network-level transcriptional patterns to human disease should be interpreted with caution.

In addition, bulk RNA sequencing inherently averages transcriptional signals across heterogeneous cellular populations. While integration with pseudobulk single-cell data partially mitigated this limitation by enabling cell-type–resolved network mapping, direct single-cell profiling of the same anatomical regions and disease stages analyzed in bulk would further strengthen cellular attribution and reduce cross-modal uncertainty.

Furthermore, the single-cell datasets used for integration were derived from multiple mouse models, brain regions, and experimental platforms. Despite standardized preprocessing and quality control, biological and technical variability across datasets may influence enrichment estimates and module–cell type associations. Future studies combining matched bulk and single-cell profiling within the same experimental cohorts will be important to refine cross-scale integration strategies.

Finally, the network associations identified in this study are correlative and do not establish causal relationships between specific cell populations, regulatory transcripts, and disease progression. Functional validation of candidate network hubs and lncRNAs, including AU020206, will be necessary to determine their mechanistic roles in amyloid-associated neuroinflammatory and transcriptional remodeling.

## Conclusions

In this study, we present an integrated multi-scale transcriptomic framework linking regional gene expression remodeling, network-level organization, and cell-type specific contributions during amyloid-driven neurodegeneration in the APP/PS1-21 mouse model. By combining bulk RNA sequencing, co-expression network analysis, and single-cell integration across cortical and hippocampal circuits, we show that Alzheimer’s disease pathology is characterized by distinct region-specific transcriptional trajectories that nevertheless converge on a conserved immune driven molecular response.

Our findings indicate that immune and inflammatory programs represent a dominant organizing feature of disease-associated transcriptional networks, while the cellular drivers of these networks differ across anatomical regions, with microglia shaping cortical responses, astrocytes strongly contributing to CA1 remodeling, and coordinated multi lineage involvement characterizing the dentate gyrus. These observations provide a framework for interpreting how region dependent cellular responses may contribute to heterogeneous disease progression.

In addition, the consistent embedding of selected lncRNAs within disease-associated networks highlights the potential contribution of regulatory noncoding transcripts to coordinated transcriptional responses that are not readily captured by single gene differential expression analyses. These findings provide a foundation for future mechanistic and translational studies aimed at dissecting immune driven network regulation and identifying potential therapeutic targets in Alzheimer’s disease.

Overall, this study provides a systems level perspective on how amyloid pathology reshapes transcriptional programs across brain regions and cell populations and establishes a foundation for future investigations into immune driven network regulation in Alzheimer’s disease.

## Data availability

The raw and processed sequencing data generated in this study will be deposited in the Gene Expression Omnibus (GEO) prior to publication. In the meantime, the data are available from the corresponding author upon reasonable request.

## Code availability

All code required to reproduce the statistical analyses and generate the supplementary tables reported in this study is publicly available at GitHub:

https://github.com/jcortessilva/APP-RNAseq-reproducibility

## Author contributions

J.A.C.S performed the bioinformatic and statistical analyses and generated all figures and supplementary tables. M.G. and S.B. performed the experimental work. T.P.C. contributed to bioinformatic analysis and supervision. F.S. contributed to figure preparation. A.F. conceived and supervised the study. All authors reviewed and approved the final manuscript

